# *In vitro* gut modeling as a tool for adaptive evolutionary engineering of *Lactiplantibacillus plantarum*

**DOI:** 10.1101/2020.10.21.349795

**Authors:** Julia Isenring, Annelies Geirnaert, Alex R. Hall, Christoph Jans, Christophe Lacroix, Marc J.A. Stevens

## Abstract

Research and marketing of probiotics demand for holistic strain improvement considering both, the biotic and abiotic gut environment. Here we aim to establish the continuous *In vitro* colonic fermentation model PolyFermS as a tool for adaptive evolutionary engineering. Immobilized fecal microbiota from adult donors were steadily cultivated up to 72 days in PolyFermS reactors, providing a long-term compositional and functional stable ecosystem akin to the donor’s gut. Inoculation of the gut microbiota with immobilized or planktonic *Lactiplantibacillus plantarum* NZ3400, a derivative of the probiotic model strain WCFS1, led to successful colonization. Whole genome sequencing of 45 recovered strains revealed mutations in 16 genes involved in signaling, metabolism, transport, and cell surface. Remarkably, mutations in LP_RS14990, LP_RS15205, and intergenic region LP_RS05100<LP_RS05095 were found in recovered strains from different adaptation experiments. Combined addition of the reference strain NZ3400 and each of those mutants to the gut microbiota resulted in increased abundance of the corresponding mutant in PolyFermS microbiota after 10 days, showing the beneficial nature of these mutations. Our data show that the PolyFermS system is a suitable technology to generate adapted mutants for colonization in colonic conditions. Analysis thereof will provide knowledge about factors involved in gut microbiota colonization and persistence.

**Importance:** Improvement of bacterial strains to specific abiotic environmental factors is broadly used to enhance strain characteristics for processing and product quality. However, there is currently no multidimensional probiotic strain improvement approach for both, abiotic and biotic factors of a colon microbiota. The continuous PolyFermS fermentation model allows stable and reproducible continuous cultivation of colonic microbiota and provides conditions akin to the host gut with high control and easy sampling. This study investigated the suitability of PolyFermS for adaptive evolutionary engineering of a probiotic model organism for lactobacilli, *Lactiplantibacillus plantarum*, to an adult human colonic microbiota. The application of PolyFermS controlled gut microbiota environment led to adaptive evolution of *L. plantarum* strains for enhanced gut colonization characteristics. This novel tool for strain improvement can be used to reveal relevant factors involved in gut microbiota colonization and develop adapted probiotic strains with enhanced functionality in the gut.

## Introduction

Microorganisms play a pivotal role in pharmaceutical, biotechnological, and food industries. The latter depends heavily on microorganisms for starter cultures, biopreservation agents, and flavor producers (1–3). Moreover, since the 1990s, there has been an increase in the production of probiotics, which are “live microorganisms that, when administered in adequate amounts, confer a health benefit on the host” (4). Strain improvement of probiotic bacteria is of major importance to meet consumer demands for functional foods and enhance competitiveness of probiotic strains. However, it demands for a multi-dimensional approach since biotic and abiotic factors are involved.

A promising solution for strain improvement is evolutionary engineering, which steers microbial evolution by exerting selective pressure (5–7). Desired mutants can be selected based on e.g. growth rate, increased survival, or retention time. This method is feasible with bacteria because short generation times and large population sizes facilitate rapid emergence and selective sweeps of mutants (8–11). It is a well-established approach to improve targeted strain characteristics like the acidification rate of *Lactococcus lactis* (12), growth of *Escherichia coli* (13) or enhanced succinate production in *Actinobacillus* and *Mannheimia* (10, 14, 15). Nonetheless, the potential of evolutionary engineering as multi-dimensional engineering within microbial consortia is not well established yet. Previously, the residence time of *Lactiplantibacillus plantarum* in the murine digestive tract was increased after repetitive administration of the longest persisting *L. plantarum* (16). However, in vivo models like mice have societal, ethical, and monetary restrictions and might therefore be substituted by *In vitro* models. Furthermore, gastrointestinal physiology and gut species composition of mice are different from humans, possibly limiting the translation (17, 18).

Continuous fermentation models are best suited for *In vitro* cultivation of gut microbiota in conditions akin to the gut (19, 20). Different PolyFermS models were successfully developed for cultivating colonic microbiota of humans of different ages and conditions, swine, murine, and chicken cecum microbiota (21–25). The continuous PolyFermS model allows testing several treatments in parallel in second-stage treatment reactors (TRs) seeded with the same gut microbiota produced in the inoculum reactor (IR) containing immobilized microbiota (21). Gut microbiota immobilization in polysaccharide gel beads leads to the maintenance of high cell density, long-term stability due to prevention of cell washout, and diversity of the simulated gut microbiota (23, 26, 27). It moreover creates a sessile bacterial fraction on the gel beads and a planktonic fraction resulting from the growth and release of sessile bacteria and further growth of planktonic cells in the bulk medium (27, 28). This mimics the gastrointestinal environment consisting of free and biofilm or mucus associated bacteria (29, 30). The PolyFermS colonic fermentation model enables operation up to several months in a highly controllable environment with multiple parameters to operate on, which is needed for evolutionary adaptation (26). We therefore hypothesized that the PolyFermS model can provide a long-term stable gut microbiota akin to the human adult colon that allows for adaptive evolution of an exogenous single strain.

In this study, we investigated the PolyFermS fermentation model as a novel tool for strain improvement via adaptive evolutionary engineering, using *L. plantarum* as model strain. *L. plantarum* originates from fermented foods (31, 32) and is detected at low levels in approximately half of healthy human gut microbiota (33). *L. plantarum* WCFS1 is a well-characterized model strain for transient probiotic lactobacilli (34, 35). A WCFS1 derivative harboring a chloramphenicol (CM) resistance gene for tracking, was cultivated in PolyFermS reactors inoculated with immobilized adult fecal microbiota for at least 100 generations. Engineered strains were phenotypically and genotypically characterized and tested for improved colonization in the PolyFermS model.

## Material and Methods

### Bacterial strains and growth conditions

Bacterial strains used during this study are listed in Supplementary Table S1. *L. plantarum* NZ3400 (36) was used as reference strain. NZ3400 is a derivative of WCFS1, harboring a CM resistance cassette (P_32_-*cat*) in a neutral locus on the chromosome. L .plantarum was grown in De Man, Rogosa and Sharpe (MRS, Labo-Life Sàrl, Pully, Switzerland) broth at 37°C, overnight. *L. plantarum* viable cells were enumerated by plating on MRS agar supplemented with CM (10 μg/ml), aerobic incubation at 37°C, overnight.

### Immobilization of adult fecal microbiota

Fecal samples were obtained from four healthy adult individuals (27-31 years old) who did not take antibiotics and probiotics for at least three months and did not show detectable microbial growth on MRS + CM plates to avoid interference with *L. plantarum* NZ3400 recovery. Out of 15 tested fecal samples, four samples that did not show microbial growth on MRS+CM and were chosen for fermentation (donor 1-4). The Ethics Committee of ETH Zürich exempted this study from review because the sample collection procedure has not been performed under conditions of intervention. Informed written consent was obtained from fecal donors. Fecal samples were immediately transferred to an anaerobic chamber within 2 h after defecation and suspended at 20% (w/v) in reduced peptone water (0.1%, pH 7; Thermo Fisher Diagnostics AG, Pratteln, Switzerland). Immobilization of 10 ml fecal slurry in polymer gel beads (gellan gum (2.5%, w/v), xanthan (0.25%, w/v) and sodium citrate (0.2%, w/v)) was performed as previously described (21, 37). The inoculum bioreactor (IR) (Sixfors, Infors, Bottmingen, Switzerland) was filled with 140 ml of vitamin supplemented MacFarlane medium (Supplementary Data S1), formulated to mimic the chyme entering the colon (20, 38), and inoculated with 60 ml of fecal gel beads. Beads were colonized in two fed-batch fermentations carried out at 37°C, pH 5.8 by controlled addition of NaOH (2.5 M), stirring at 180 rpm, and replacing 100 ml of spent medium with fresh medium after 16 h (21). Anaerobiosis was set by purging the bioreactor headspace with CO2 and monitored by redox potential probes.

### Proximal colon fermentation in the PolyFermS model using immobilized human gut microbiota

Continuous proximal colonic *In vitro* fermentations with immobilized human adult gut microbiota were performed in bioreactors as reported previously (26, 39). All fermentations were operated to mimic the adult proximal colon as described above for batch fermentation. Fresh medium, was continuously added to the IR (25 ml/h) and fermented medium was removed to maintain a working volume of 200 ml. Because short-chain fatty acids (SCFAs) are the main fermentation end products of the gut microbiota, their stable production is a convenient, easily measurable marker for stability of continuous intestinal fermentation models (21, 40). The IR was run in continuous mode for at least 10 days to reach metabolic stability indicated by lower than 10% day-to-day variation (25, 40) in SCFAs concentration, before connecting second-stage TRs. TRs were inoculated by the IR at 1.25 ml/h and fed at 23.75 ml/h. TRs were operated continuously for four days prior to *L. plantarum* supplementation in order to establish a gut microbiota activity akin to the IR. Addition of planktonic or immobilized *L. plantarum* was tested in IR and TRs with different donor microbiota (Fig. 1). In case of two successive treatment periods in the same reactor, the reactor was disconnected after the first treatment period, sterilized, re-connected and stabilized for four days before starting a second treatment.

**Fig. 1:**
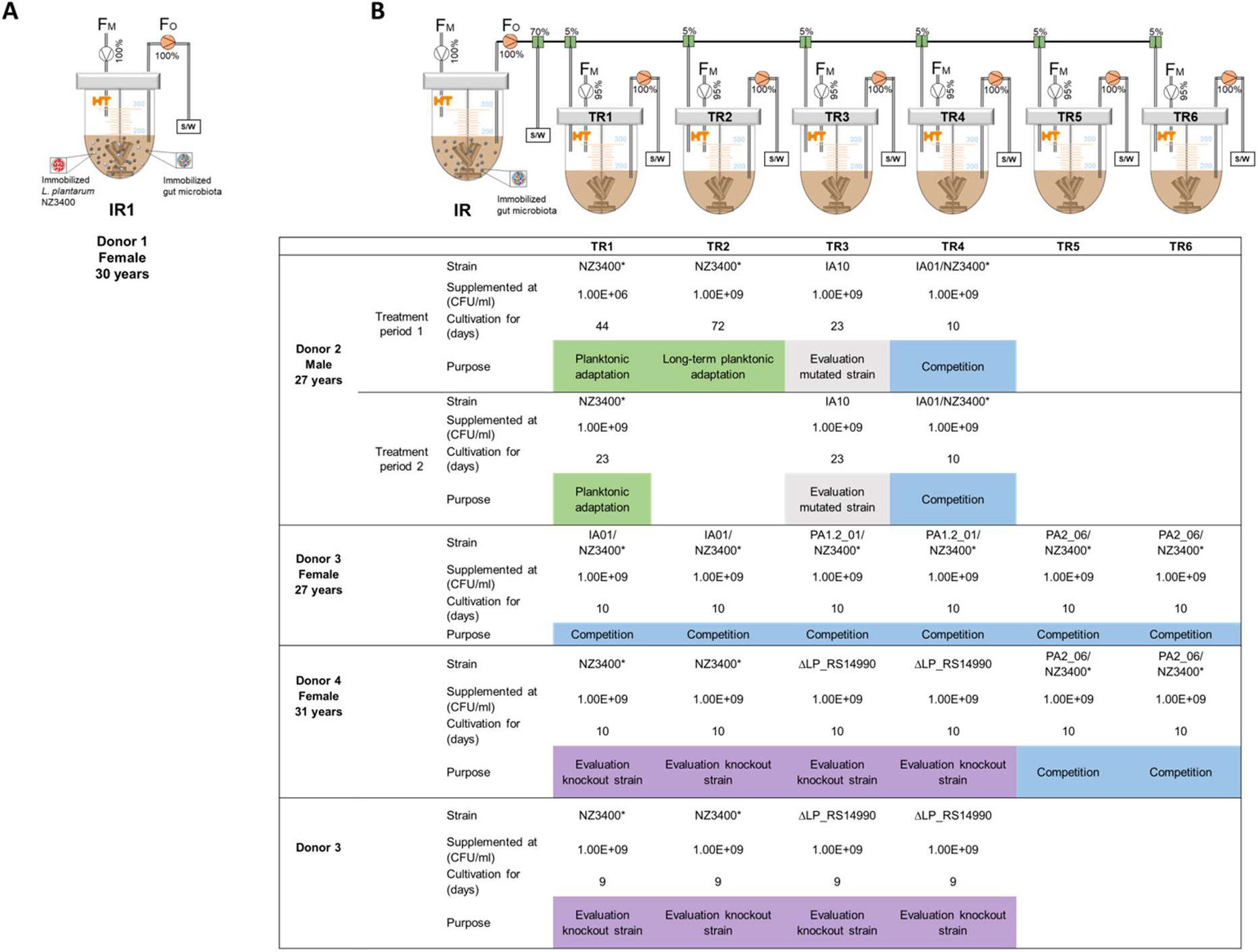
PolyFermS setup of immobilized and planktonic adaptation and further experiments in *In vitro* human adult gut microbiota. A) Immobilized *L. plantarum* NZ3400 was added to immobilized fecal gut microbiota in IR1 from donor 1 during a single-stage fermentation. B) Immobilized fecal microbiota of donors 2-4 were cultivated in the inoculum reactor (IR), which was used to inoculate second stage reactors (TRs) that were supplemented with *L. plantarum* after stabilization. NZ3400: reference strain; NZ3400*: new stock from single colony isolate of NZ3400; IA10, IA01, PA1.2_01, PA2_06: recovered *L. plantarum* mutants; *L. plantarum* ΔLP_RS14990: LP_RS14990 gene deletion strain. FM: Inflow MacFarlane medium, FO: Reactor outflow S/W: Sampling/Waste.

Reactor effluent samples were taken daily to monitor the fermentation process, centrifuged for 10 min at 14 000 g, 4°C and stored at −20°C. Pellets were used for DNA extraction and supernatants for metabolite analysis.

### Adaptive evolutionary engineering of *L. plantarum*

#### Adaptive evolution using immobilized *L. plantarum* NZ3400

Adaptive evolution of immobilized *L. plantarum* NZ3400 was tested in a continuously run single-stage IR (IR1) inoculated with donor 1 fecal beads (Fig. 1, A). A *L. plantarum* overnight culture was harvested at 4°C, 4000 g for 10 min and washed twice in Phosphate-Buffered Saline (PBS), pH 6.2. Immobilization of *L. plantarum* was done as described for fecal samples under aerobic conditions. Beads were colonized during two pH-controlled batch cultures at 37°C for 16 h with stirring at 150 rpm. Colonized beads were washed in PBS, supplemented with cryoprotective buffer (41) and stored at - 80°C. Before use, *L. plantarum* beads were reactivated during two batch cultures as described for colonization and washed twice in PBS. *L. plantarum* viable cell counts were determined by crushing 1 g of beads in PBS with a spatula and plating serial dilutions. Four g of beads (5*10^9^ colony forming units (CFU) *L. plantarum*/g) were added to the single-stage IR1 and cultivated for 53 days.

#### Adaptive evolution using planktonic *L. plantarum* NZ3400*

Due to microbial growth on MRS+CM plates of donor 1 fecal sample at this time, a new donor was chosen for the planktonic adaptation. To prevent carry-over of possible mutants present in the initial NZ3400 stock culture, a new stock from a single NZ3400 colony isolate, designated NZ3400*, was produced. NZ3400* was subjected to PacBio sequencing and used for all subsequent planktonic supplementation trials. NZ3400 and NZ3400* differed in eight SNPs but none thereof in SNP affected genes of recovered *L. plantarum*. Adaptation of planktonic *L. plantarum* NZ3400* was tested in TRs continuously inoculated by the effluent from IR2 containing beads with immobilized fecal microbiota of donor 2 (Fig. 1, B). Because the PolyFermS model was built with six TRs, two consecutive treatment periods were performed to test all treatments. *L. plantarum* strains were grown overnight, harvested, washed twice in PBS, re-suspended in MacFarlane medium and added to the TRs to a final level of 10^9^ CFU *L. plantarum*/ml effluent. Long-term planktonic adaptation was tested in TR2 operated for 72 days and repeated in TR1 (period 1) and TR1 (period 2) for 44 and 23 days, respectively (Fig. 1, B). Furthermore, the strain IA10, recovered from the immobilized adaptation, was added in planktonic state to TR3 (period 1) and TR3 (period 2) containing donor 2 gut microbiota to investigate effects of adaptations that occurred during the immobilized adaptation (Fig. 1, B).

### Metabolite analysis of the continuous colon fermentation

Reactors were sampled daily for analysis of the SCFAs acetate, butyrate and propionate, branched-chain fatty acids isobutyrate, isovalerate and valerate, and intermediate metabolites lactate and formate (42). Concentrations were determined by high-performance liquid chromatography (HPLC) as described previously (23).

### Microbial profiling by 16S rRNA gene amplicon sequencing

Genomic DNA of fecal and effluent samples was extracted using the FastDNA^®^ SPIN Kit for Soil (MP Biomedicals, Illkirch, France) according to the manufacturer’s instructions. The V4 region of the 16S rRNA gene was amplified with the primers 806R (5′-GGACTACHVGGGTWTCTAAT-3′) and 515F (5′-GTGCCAGCMGCCGCGGTAA-3′). Amplicons were barcoded PCR-based. Library preparation and sequencing (Illumina, CA, USA) using an Illumina MiSeq flow cell with V2 reagent kit for 2 × 250-bp paired-end Nextera chemistry supplemented with 10% of PhiX was performed in collaboration with the Genetic Diversity Center (GDC, ETH Zürich, Switzerland).

Raw data obtained from 16S rRNA sequencing were processed using Cutadapt (43) and DADA2 pipeline (44) to obtain amplicon sequence variants. Taxonomy was assigned using SILVA database (v.132) (45) (full method in Supplementary Data S1).

### Recovery of *L. plantarum* from the gut microbiota

*L. plantarum* colonization was determined by plating on MRS + CM agar. Combination of the MRS selectivity for lactobacilli and enterococci together with aerobe incubation and presence of antibiotics allowed growth repression of all other bacteria. Data were compared to the theoretical washout curve determined for absence of growth, from the following equation: c_t_=c_0_*e^(t/RT)^ (39), where c_0_ and c_t_ represent cell concentration at time point 0 and t, respectively, and RT corresponds to the retention time. Colonies were randomly picked, incubated in MRS + CM overnight, mixed 1:1 with 60% (v/v) glycerol (Sigma-Aldrich Chemie GmbHm Buchs, Switzerland) and stored at −80°C.

Natural biofilm formed in TR2, used for long-term planktonic adaptation, and the repetition experiment in TR1 (period 1). To recover *L. plantarum* from biofilms, the vessels were emptied and washed twice with PBS. Remaining biofilm was removed and homogenized with glass beads (5 mm, VWR International AG, Dietikon, Switzerland) in dilution solution containing 0.85% NaCl and 0.1% peptone from casein (w/v, VWR International AG, Dietikon, Switzerland). Dilutions were plated with subsequent strain recovery and storage performed as described above.

### Phenotypic characterization of recovered *L. plantarum* strains

Growth behavior of recovered *L. plantarum* strains was analyzed in MRS supplemented with each of the main SCFAs of the human gut microbiota, acetate (50, 75, 100 mM), butyrate (15, 30, and 45 mM) and propionate (15, 30, and 45 mM) in similar concentrations as measured during colonic fermentations (see Fig. S2 in the supplemental material). The abiotic gut fermentation environment was simulated in Effluent-MacFarlane-Sugar (EMS) medium consisting of filter-sterilized PolyFermS effluent, MacFarlane medium in a ratio 9:1 and 0.75% (w/v) glucose (see Fig. S1 in the supplemental material). Glucose was added since *L. plantarum* was unable to grow in MacFarlane medium. For comparison of the effect on adaption in immobilized and long-term planktonic adaptation trials performed in TR2 (Fig. 1, B), recovered strains were divided into four groups based on their origin of isolation: (1) 11 strains from the effluent at the late stage of immobilized adaptation after 53 days, (2) 14 strains from the effluent during day 7 and day 23 (early planktonic adaptation), (3) 19 strains from the effluent during day 60 and day 72 (late planktonic adaptation) and (4) 25 strains from the biofilm of planktonic adaptation after 72 days. Strains were isolated at an early stage after seven (stable *L. plantarum* colonization) and 23 days and late stage of 60 and 72 days to increase the chance to observe adaptation. Biofilm was sampled on the last day of fermentation because the reactor had to be emptied for biofilm sampling. Growth analysis was done in 96-well tissue culture test plates (Bioswisstec AG, Schaffhausen, Switzerland). Wells were filled with 200 μl of medium and inoculated with the potentially adapted *L. plantarum* at 37°C. Growth was monitored by optical density (OD) measurement at 600 nm in a plate reader after 24 h (PowerWaveTMXS, Bio-Tek Instrument Inc., Winooski, VT, US) in biological triplicates.

Phenotype stability was assessed by repeated daily culturing of strains in MRS for 28 days, approximately 190 generations, as presented above. Stability was measured after 1, 7, 14, 21 and 28 days, which corresponds to the time of transcriptome homogenization among *L. plantarum* strains isolated from different habitats (46).

### Complete genome sequencing and data analysis

*L. plantarum* genomic DNA was isolated via lysozyme-based cell lysis (47) followed by purification using the Wizard^^®^^ Genomic DNA Purification Kit according to the manufacturer’s instructions (Promega, Dübendorf, Switzerland). The genome of the reference strain NZ3400* was sequenced at the Functional Genomics Center Zurich (Zürich, Switzerland) on PacBio RS II (Pacific Biosciences, Menlo Park, CA, USA) using one SMRT cell. Reads were assembled using HGAP assembly as described previously (48). All other *L. plantarum* strains were sequenced using Illumina Miniseq (Illumina, CA, US) with 250 bp paired-end reads at the Institute for Food Safety and Hygiene, University of Zurich (48). Reads were merged and mapped to the reference genome *L. plantarum* NZ3400* using CLC Genomic Workbench 11.0 (Qiagen CLC bio, Aarhus, Denmark) using default parameters. Single nucleotide polymorphisms (SNPs) were extracted using Parsnp (49). SNPs were confirmed by Sanger Sequencing (Eurofins GATC, Biotech GmbH, Konstanz, Germany). SNP related changes in amino acid sequence were determined in CLC Genomic Workbench.

### Competition experiments in the human gut microbiota

Competition experiments between NZ3400* and the mutant strains *L. plantarum* IA01, PA1.2_01 and PA2_06 were performed to determine the effect of the mutations on gut microbiota colonization (Fig. 1, B). NZ3400* was paired with each of the three mutant strains in a 1:1 ratio to reach 10^9^ CFU/ml reactor effluent to the gut microbiota and cultivated for ten days. Ten days was sufficient to obtain stable *L. plantarum* colonization for at least four days but also as short as possible to minimize the chance of proliferation of new mutants. Relative abundance of the strains in the complex community was determined by measuring allele frequency of the genes LP_RS14990 and LP_RS15205, respectively, via Pyrosequencing (full method in Supplementary Data S1).

### Stability of mutations under standard cultivation conditions

Stability of the mutations in *L. plantarum* IA01 and PA2_06 was investigated during repeated daily cultures in MRS for 12 days, approximately 81 generations, performed in triplicates. Cultivation of NZ3400* served as control. Allele frequency was determined via Pyrosequencing as described above.

### Plasmid construction and LP_RS14990 gene replacement of *L. plantarum* NZ3400*

Six mutants carrying the mutation in LP_RS14990 were recovered from independent adaptation experiments. To investigate the involvement of this gene in gut microbiota colonization, a *L. plantarum* ΔLP_RS14990 knockout was constructed by double-crossover gene replacement in *L. plantarum* NZ3400* (full method in Supplementary Data S1). The knockout strain was subsequently added to the gut microbiota and colonization levels were compared to the reference strain NZ3400* (Fig. 1, B).

### In silico analysis of LP_RS15205 in *L. plantarum*

LP_RS15205 was affected by identical mutation in four strains recovered from two independent adaptation experiments. Since this mutation was also found in the background gut microbiota, in silico analysis of this locus was performed. Complete genome sequences of Firmicutes (n = 557), Bacteroidetes (n = 218), Actinomycetes (n = 457) and γ-Proteobacteria (n = 638) from the NCBI genome database were downloaded in May 2020. The amino acid sequence of LP_RS15205 was blasted against these genomes using standard settings and significant hits were aligned using MUSCLE (50).

### Data analysis

Statistics for growth experiments were calculated in R (version 3.6.2) using one-sample t-test for comparison to *L. plantarum* NZ3400* and paired sample t-test for comparison within recovered *L. plantarum* groups. Values represent mean values ± standard deviation. The heatmap was generated using the R pheatmap package and Euclidean distance measure. Allele frequency determination by Pyrosequencing was calculated based on three extracted DNA samples of the same time point. Graphs were created using GraphPad Prism^®^ version 8 (GraphPad Software Inc., San Diego, CA, USA).

## Results

### PolyFermS model allows stable cultivation of adult gut microbiota

The PolyFermS model operated to mimic the adult proximal colon was used to provide a gut microbiota environment for evolutionary adaptation of *L. plantarum* NZ3400. Adaptation of immobilized *L. plantarum* was performed in IR1 and adaptation of planktonic *L. plantarum* was tested in TRs continuously inoculated with IR2 microbiota (Fig. 1). Metabolic stability of IR1 (see Fig. S2, A in the supplemental material) was achieved after one week with main SCFAs acetate, propionate and butyrate at 73 ± 7, 21 ± 4 and 19 ± 3 mM over two month fermentation, respectively. IR1 microbiota was dominated by Firmicutes and Bacteroidetes accounting for 44 ± 5 and 47 ± 2% of the total population during days 44-46, respectively, 48 ± 7 and 41 ± 2% during days 57-59, respectively, and 52 ± 4 and 36 ± 4% during days 61-65, respectively (see Fig. S3, A in the supplemental material). This stability was also observed on family level (see Fig. S3, B in the supplemental material). IR2 microbiota reached metabolic stability (see Fig. S2, B in the supplemental material) after one week with the main SCFAs acetate, propionate and butyrate at 84 ± 5, 32 ± 8 and 23 ± 4 mM, respectively, during two months of culture. IR2 had a different microbiota composition from IR1 (see Fig. S3, C&D in the supplemental material). Bacteroidetes dominated the gut microbiota compared to Firmicutes with 63 ± 1% and 36 ± 1% during days 24-26, respectively, 68 ± 4 and 30 ± 4% during days 41-43, respectively, and 66 ± 1 and 30 ± 1% during days 89-91, respectively (see Fig. S3, C in the supplemental material). Stability was maintained up to 90 days on family level (see Fig. S3, D in the supplemental material). Microbiota composition of the IR was successfully transferred to and maintained in the TRs (see Fig. S3, D in the supplemental material). Both, metabolic activity (see Fig. S2, C-E in the supplemental material) and composition were reproduced in the TRs.

### Prolonged cultivation of *L. plantarum* in *in vitro* human gut microbiota

To investigate the potential of the continuous *in vitro* gut fermentation model PolyFermS for evolutionary engineering, immobilized *L. plantarum* NZ3400 was added to the stabilized microbiota in IR1 at an initial concentration of 10^8^ CFU/ml effluent. NZ3400 decreased at the rate of the theoretical wash-out during the first four days (Fig. 2, A), followed by colonization between 10^2^ CFU/ml and 10^4^ CFU/ml during the 50-days fermentation. *L. plantarum* was able to maintain a self-sustaining population, an observation which will be referred to as colonization.

**Fig. 2:**
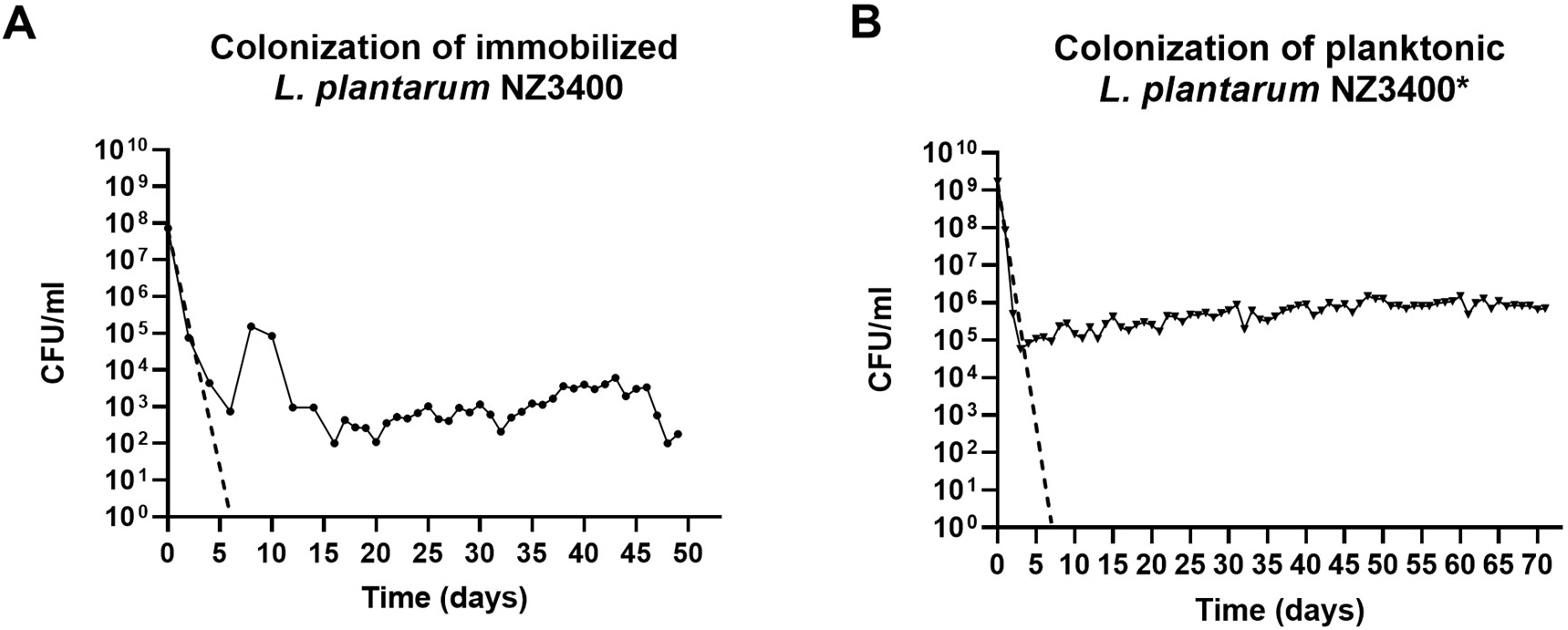
Colonization levels of *L. plantarum* during immobilized and planktonic adaptation in adult gut microbiota. A) Immobilized *L. plantarum* NZ3400 (four g of colonized beads containing 5*10^9^ CFU *L. plantarum*/g) was added to the gut microbiota (●) and B) NZ3400* at 10^9^ CFU/ml effluent in planktonic state (■). Dashed line indicates the theoretical wash-out of the system. Cell count is depicted on the y-axis and days of cultivation in the microbiota on the x-axis, whereas 0 corresponds to the day of *L. plantarum* supplementation.

The PolyFermS model was further evaluated for long-term adaptation of planktonic *L. plantarum* NZ3400* in TR2, fed by donor 2 gut microbiota. Cell counts decreased from 10^9^ to 10^5^ CFU/ml during the first four days after spiking (Fig. 2, B), at the rate of the wash-out. Thereafter, colonization steadily increased to 10^6^ CFU/ml during 72 days. Repeated supplementation of *L. plantarum* with 10^6^ and 10^9^ CFU/ml in TR1 (period1) and TR1 (period2) resulted in stable colonization at different levels of 10^6^ and 3*10^4^ CFU/ml, respectively. Therefore, *L. plantarum* colonization at a donor specific level was demonstrated for more than 50 days and 150 generations.

### Recovered *L. plantarum* strains are phenotypically adapted to the gut microbiota environment

To test for *L. plantarum* adaptation during gut microbiota cultivation, recovered strains were grown in SCFA concentrations comparable to those in the gut fermenter. Average growth of strains from immobilized adaptation measured by OD was impaired in MRS and MRS supplemented with 30 mM propionate or butyrate compared to the reference strain NZ3400* (Table 1). Strains from early stage of long-term planktonic adaptation behaved similarly to the reference NZ3400* in all tested media. However, strains from late planktonic adaptation and biofilm grew better in MRS plus acetate (+0.11 and +0.07, respectively), less in standard MRS (−0.14 and −0.11, respectively), and similar in MRS plus propionate or butyrate (Table 1) compared to NZ3400*. Moreover, strains from biofilm and late planktonic adaptation grew significantly better in the reactor-effluent-mimicking EMS medium compared to strains from early adaptation (Table 1).

**Table 1:**
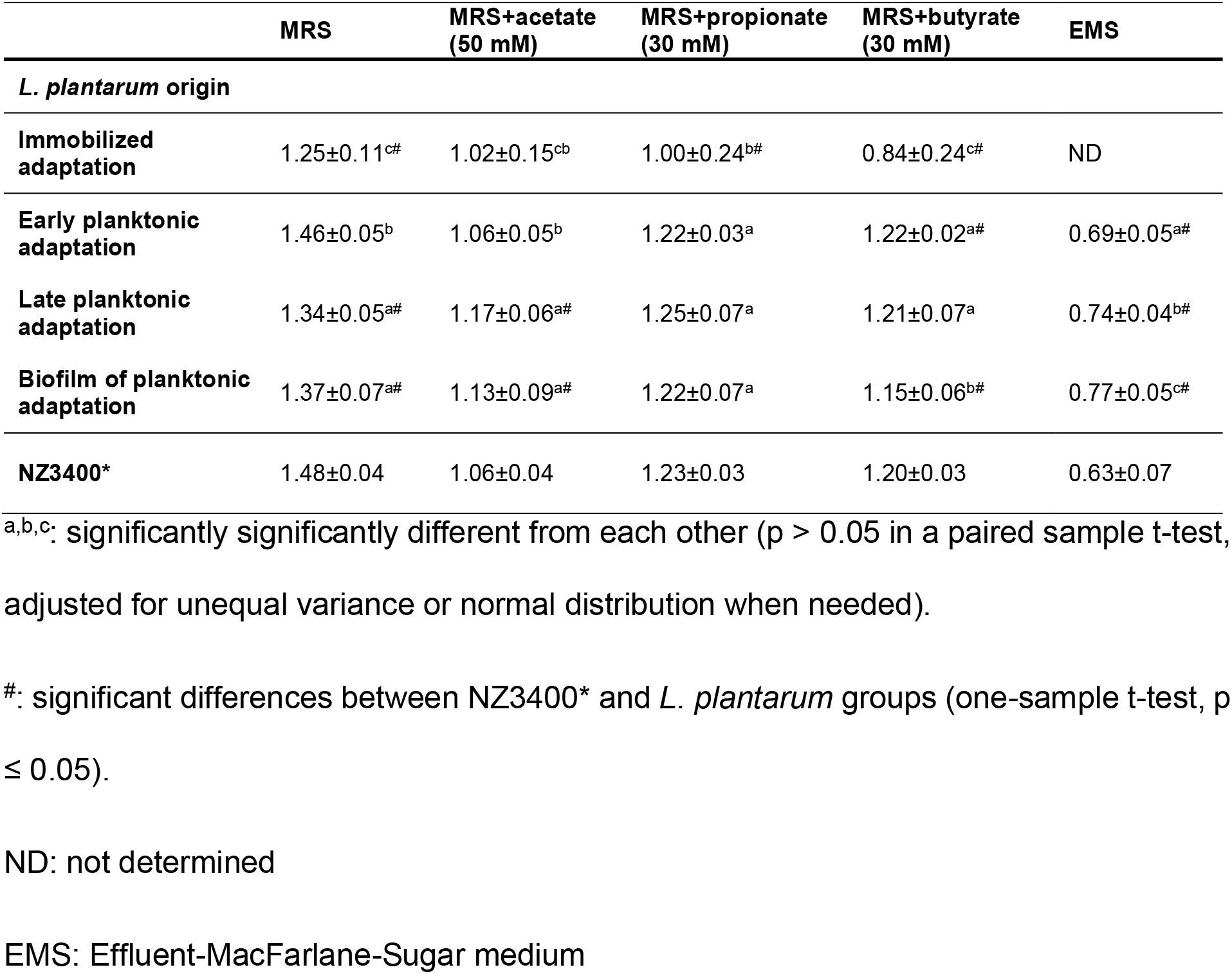
Growth of recovered *L. plantarum* strains in different media. The values reported are OD_600nm_ after 24 hours. Values obtained for NZ3400* represent mean ± standard deviation of biological triplicates. All other values represent mean ± standard deviation of all strains recovered from one adaption period, whereas each recovered strain was measured in biological triplicates. Immobilized adaptation: n= 11 strains, early planktonic adaptation: n= 14 strains, late planktonic adaptation: n= 19 strains, biofilm of planktonic adaptation: n= 25 strains.

When clustered according to growth performance in MRS and MRS supplemented with acetate (50 mM), butyrate (30 mM) and propionate (30 mM), strains from immobilized and planktonic adaptation were clearly separated (Fig. 3). Further, strains recovered from immobilized adaptation exhibited a higher growth variability than strains isolated from the long-term planktonic adaptation (see Fig. S4 in the supplemental material). The reference NZ3400* clustered with strains isolated from early planktonic adaptation and clearly separate from strains isolated from effluent and biofilm at the end of adaptation (Fig. 3). Altogether, these results strongly hint towards adaptation of *L. plantarum* during prolonged cultivation in the gut microbiota. Furthermore, strains did not cluster according to the reactor they were isolated from, suggesting that the adaptation pattern is not dependent on reactor but rather time point of isolation.

**Fig. 3:**
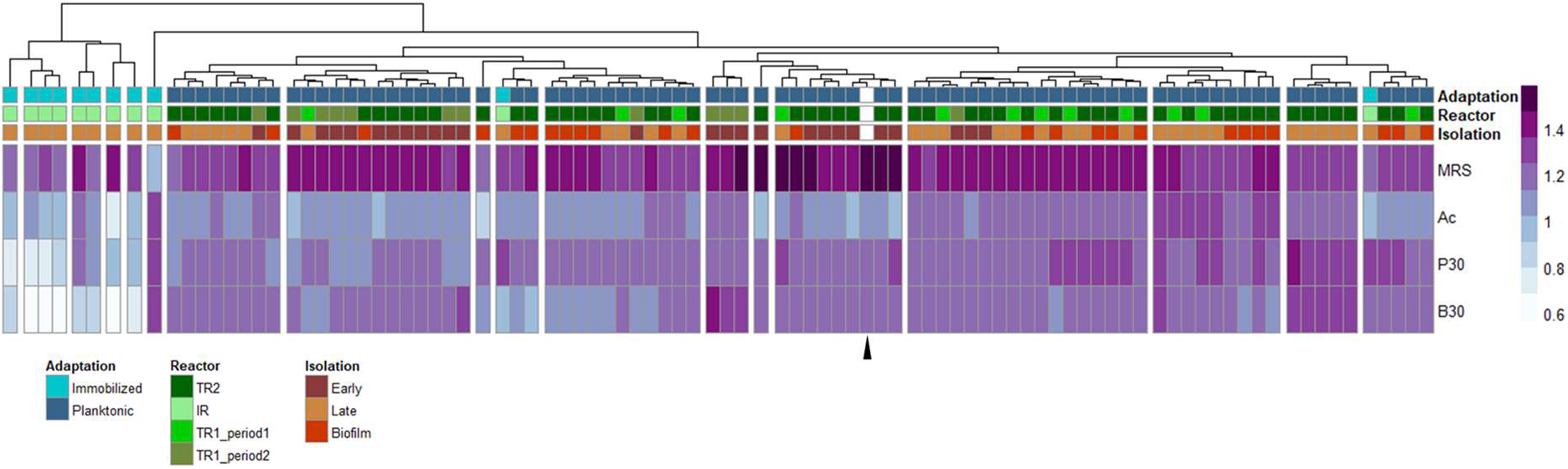
Heatmap visualization of growth behavior of potential mutant *L. plantarum* strains versus their reference strain NZ3400* in modified MRS medium (single combination of MRS with acetate (50 mM), butyrate (30 mM) or propionate (30 mM)). Image symbols: (i) Adaptation: indicates strain origin from immobilized or planktonic adaptation trials. (ii) Reactor: describes the general experiment type for long-term planktonic adaptation (TR2) and the repetition of planktonic adaptation in TR1 (period 1) and TR1 (period 2). (iii) Isolation: indicates biofilm/time point of isolation (early or late) of a strain during the adaptation cycles. Heatmap colors stand for measured ΔOD_600nm_ values after 24h. The black triangle represents growth of *L. plantarum* NZ3400*. Ac: acetate, P: propionate, B: butyrate.

Phenotypes of these adaptations were stable for at least 190 generations (see Fig. S5 in the supplemental material). This strongly suggests that observed phenotypes are caused by mutations rather than physiological variations.

### Mutations in adapted *L. plantarum* strains hint towards adaptive evolution

Stable altered phenotypes of recovered *L. plantarum* strains strongly suggest that these strains harbor mutations. Therefore, whole-genome sequencing of 45 strains randomly selected from adaptations experiments was performed. Comparison to the reference genome NZ3400* revealed 15 strains without any genotypic differences. Out of 18 SNPs confirmed by Sanger sequencing, two were detected in non-coding regions. The remaining 16 SNPs were found in genes involved in signaling, metabolism, transport, and cell surface (Table 2).

**Table 2:**
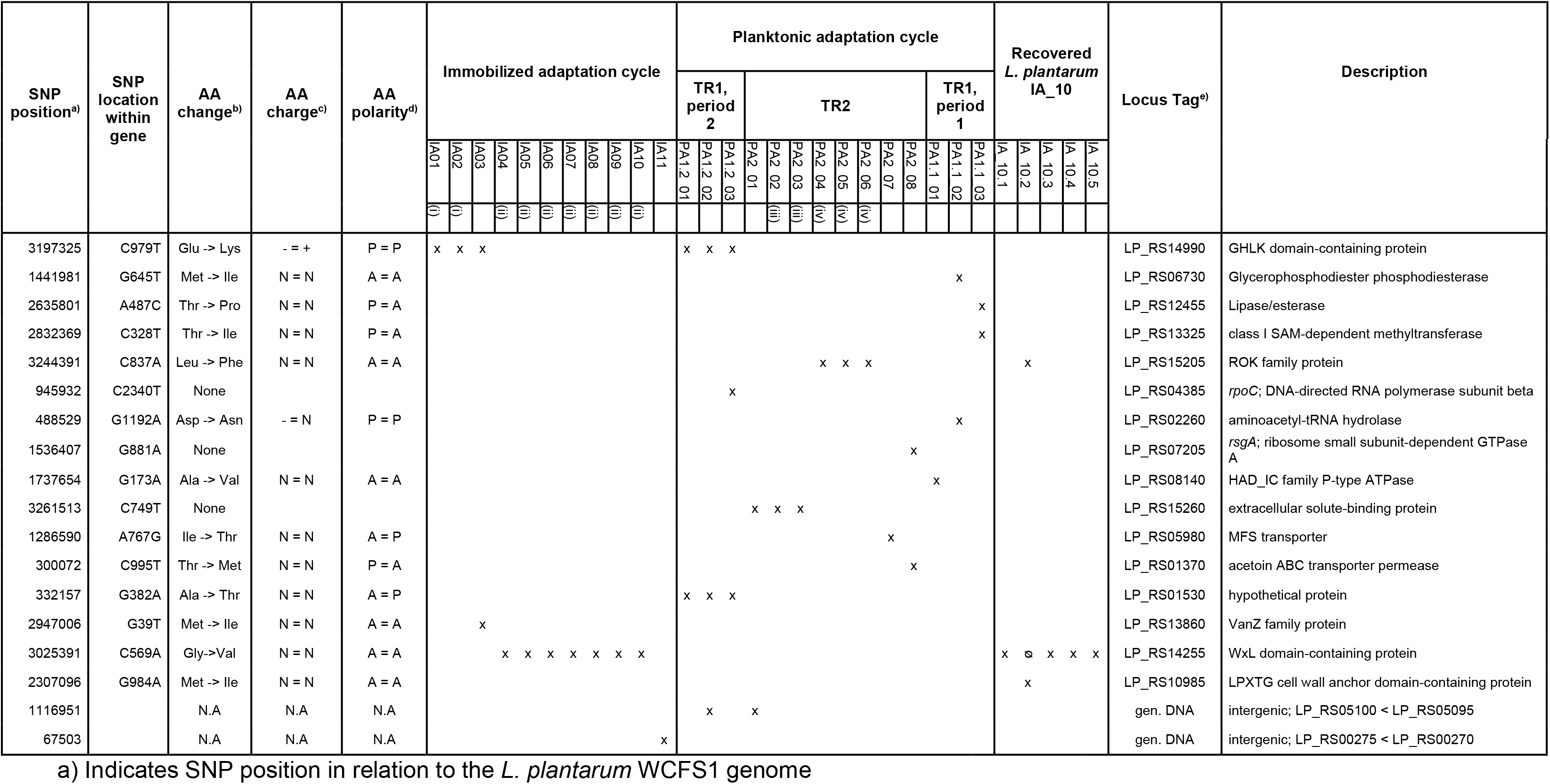
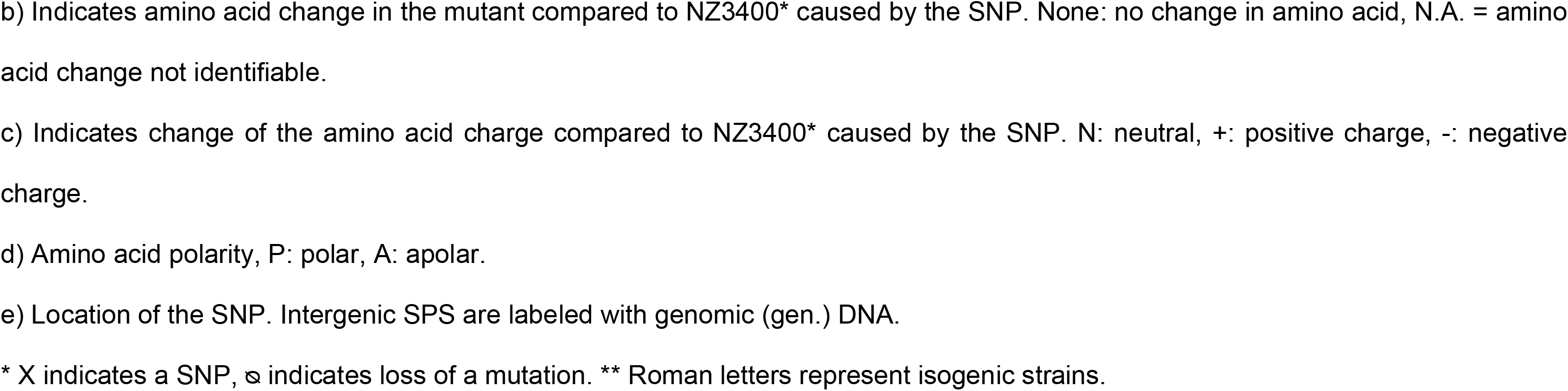
Identified genotypic changes of recovered *L. plantarum* strains compared to NZ3400*.

The 11 mutated of 12 sequenced strains from the immobilized adaptation belonged to four different genotypes. Among these 11 strains, a mutation in the cell surface protein gene encoded by LP_RS14255 was found seven times. Further, a mutation in LP_RS14990, encoding a histidine kinase domain, was found thrice (Table 2). Eight of 15 *L. plantarum* strains that were recovered at late stage of the long-term planktonic adaptation were mutated, resulting in five different genotypes (Table 2). Strikingly, a SNP in LP_RS14990 occurred independently during planktonic and immobilized adaptation. Moreover, the SNP in LP_RS15205 and in the intergenic region between LP_RS05100 and LP_RS05095 were found in strains isolated from two different reactors.

Immobilized adaptation resulted in a bigger fraction of mutated strains but consisted predominantly of two isogenic lineages. Planktonic adaptation resulted in less frequent mutagenesis, yet higher mutant diversity. This suggests a difference in adaptation pressure, as already observed for the phenotypic screening. Recovery of some identical mutants from different adaptation experiments suggests that some of the observed mutations are involved in adaptation to the gut microbiota.

### Mutations in LP_RS14990 and LP_RS15205 are beneficial for *in vitro* gut microbiota colonization

The mutation in the histidine kinase protein gene LP_RS14990 and the ROK protein gene LP_RS15205 occurred independently more than once in adaptation experiments. We therefore tested the fitness of each of the mutants *L. plantarum* PA2_06 (C837A in LP_RS15205), IA01 (C979T in LP_RS14990) or PA1.2_01 (C979T in LP_RS14990 and C837A in LP_RS15205) in competition experiments with the reference strain NZ3400*. Ten days of cultivation resulted in an increased abundance of all tested mutants in the gut microbiota when compared to NZ3400* (Table 3).

**Table 3:**
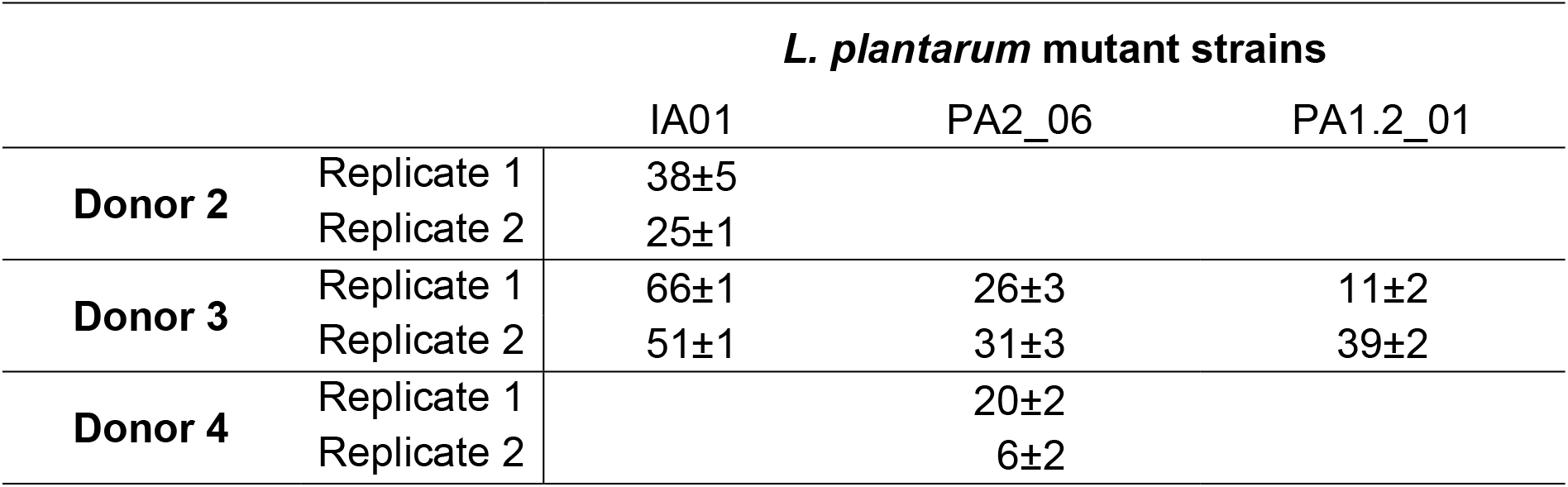
Increase in relative abundance (in %) of *L. plantarum* mutants during 10 days of competition against NZ3400* in *In vitro* human gut microbiota. Presented values show the increase of the ratio of *L. plantarum* mutant to the reference strain NZ3400* after 10 days of cultivation compared to the time point of inoculation. Values represent mean ± standard deviation of three DNA samples isolated at the same time point from the same reactor.

Pyrosequencing indicated that donor 2 gut microbiota had a *L. plantarum* background. Remarkably, the Pyrogram of this background at the position LP_RS15205 C837A was identical to the Pyrogram of a sample containing NZ3400* and PA2_06. This shows that the nucleotide variation of both, NZ3400* and PA2_06, also occurs naturally (see Fig. S6 in the supplemental material).

### Mutation C979T in LP_RS14990 is stable under standard culturing conditions

After observing increased fitness of mutants compared to the reference strain, it was investigated whether the mutations of *L. plantarum* IA01 in LP_RS14990 and *L. plantarum* PA2_06 in LP_RS15205 are stable during daily repeated batch cultures without the adaptation pressure of the gut microbiota. Stability of the LP_RS14990 mutation in the IA01 strain was observed during 12 batch cultures. However, the mutation C837A in strain PA2_06 was not stable since the reference strain nucleotide reoccurred at 3.5 ± 0.15% after 12 days (see Table S3 in the supplemental material). Investigation of NZ3400* in repeated MRS batch cultures revealed no occurrence of the SNPs of the mutants.

### LP_RS14990 gene replacement in *L. plantarum* NZ3400* results in delay of gut microbiota colonization

To investigate the role of the LP_RS14990 gene in gut microbiota colonization, a ΔLP_RS14990 gene replacement strain was constructed and its colonization compared to NZ3400*. NZ3400* started to colonize the gut microbiota of donor 3 on day one and donor 4 at day three since levels were above the wash-out curve (Fig. 4). Levels of ΔLP_RS14990 in both gut microbiota decreased more rapidly than the wash-out curve until day three, suggesting cell death (Fig. 4). Strain ΔLP_RS14990 started to colonize the gut microbiota of both donors only after four days, later than the reference strain.

**Fig. 4:**
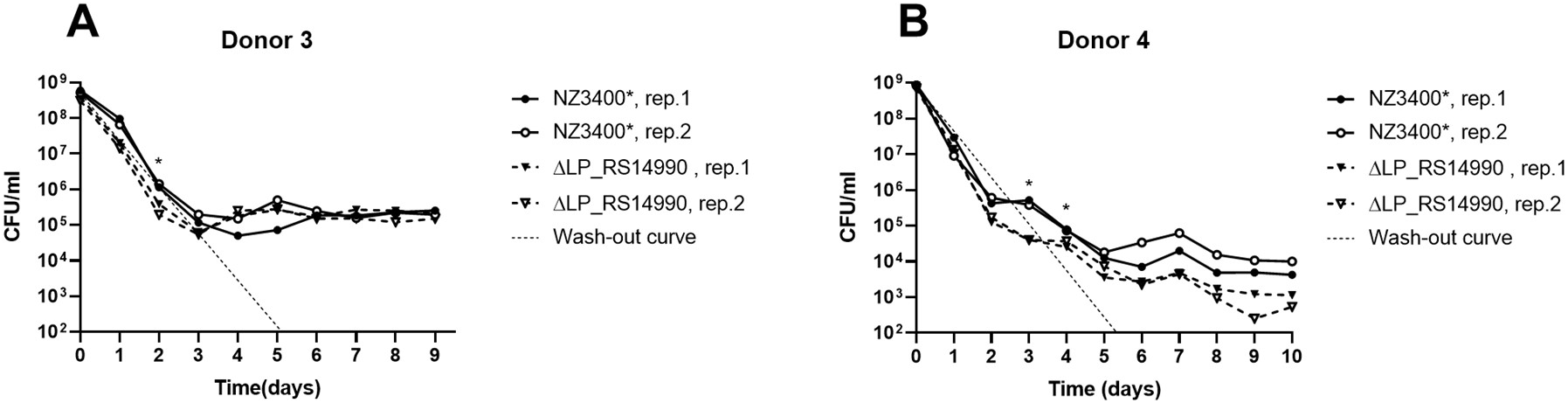
Colonization of ΔLP_RS14990 compared to the reference strain NZ3400*. *L. plantarum* strains were added to a level of 10^9^ CFU/ml reactor effluent. A) *L. plantarum* ΔLP_RS14990 and NZ3400* were added into each of the two reactors (rep.1, rep.2) in donor 3 gut microbiota. B) *L. plantarum* ΔLP_RS14990 and NZ3400* were added into each of the two reactors (rep.1, rep.2) in donor 4 gut microbiota. Dotted lines indicate the theoretical wash-out curve of the system. Cell count is depicted on the y-axis and days of cultivation in the microbiota on the x-axis, whereas 0 corresponds to the day of *L. plantarum* supplementation. *: significantly different colonization levels between NZ3400* and ΔLP_RS14990 (p ≤ 0.05, paired sample t-test).

### In silico analysis of LP_RS15205 in *L. plantarum*

Occurrence of the SNP C837A (L279F) in LP_RS15205 in *L. plantarum* strains from different reactors and in the background of the gut microbiota suggests an important function of this SNP in survival of *L. plantarum* in the gut microbiota. In silico analysis of LP_RS15205 revealed that C837A lies in a conserved domain with an ANLxGA motive that is found in Firmicutes and γ-Proteobacteria (data not shown). The function of this domain is unknown, but its conservation throughout two phyla strongly suggests that it might be important for protein function and the mutation ANLxGA to ANFxGA might have an impact on protein functionality.

## Discussion

Strain improvement is of major importance for food industries and probiotic cultures. Classical evolutionary adaptation approaches focus on specific strain characteristics in abiotic environments like improved performance of starter cultures or lactic acid bacteria (51, 52). However, for probiotics, improvement of gut colonization needs to consider abiotic and biotic factors. Here we examined the suitability of the *in vitro* gut fermentation model PolyFermS to provide a gut microbiota environment for evolutionary engineering of *L. plantarum* NZ3400 towards a human adult gut microbiota. Immobilization of the donor gut microbiota and cultivation in the PolyFermS allowed reproduction of distinct gut microbiota representative for human adults (53, 54). The achieved long-term metabolic and compositional stability allowed the generation of adapted mutants and the use of several donors increased the external validity of results on observed adaptations. *L. plantarum* supplementation method and donor microbiota seem to influence phenotypic and genotypic adaptations.

Even though observed phenotypic differences of recovered *L. plantarum* were small, *L. plantarum* strains recovered from the immobilized adaptation showed very limited adaptation in contrast to recovered strains from planktonic adaptation trials, although abiotic conditions were highly similar. It thus is assumed that differences in adaptation patterns are caused by the supplementation method. Immobilization protects against stress and entraps cells physically in the system, leading to decreased adaptation pressure (55–57), which is in accordance with our phenotypic screening where strains from immobilized adaptation showed less adaptation. Polymer beads create mucosal-like attachment sites (58) and growth in beads results in a gradual release of sessile cells from the surface that then grow as planktonic cells in the bulk medium (37). The result is a mixed population consisting of cell lineages undergoing varying number of growth cycles in the planktonic state. This might explain the more diverse phenotypic adaptation pattern among strains from the immobilized adaptation. It was further shown that immobilized cells are genetically more stable than planktonic cells (59–62), which was observed in our study where recovered strains from immobilized adaptation mainly consisted of two isogenic lineages.

Two observations suggest that the observed genotypic changes in isolated strains were involved in adaptive evolution. The first is the known function of several of the genes affected by SNPs. Activated genes of *L. plantarum* in the murine digestive tract were involved in carbohydrate transport, metabolism and cell surface, and sugar-related functions, molecule biosynthesis and stress response (63, 64). Further, exposure of *L. plantarum* to the murine intestinal tract predominantly resulted in mutations in genes encoding cell wall-associated proteins (16). The SNPs identified in our study belong to the categories mentioned above. Remarkably, the latter study identified a SNP in a glycerophosphodiester phosphodiesterase encoding gene, a gene also affected by a SNP in a recovered strain in our study. In summary, the function of mutated genes are in agreement with previously observed responses of *L. plantarum* to the in vivo intestinal environment. This shows similar selection pressures in the PolyFermS system compared to in vivo settings.

The second observation indicating adaptive evolution in our experiment is that three mutations were found independently in multiple experiments. Since mutations occur continuously, −80°C stock cultures are rarely isogenic and the mutants found in our study could have been already present as subculture in the NZ3400 stock (16). To counteract this bias, two different stock cultures, NZ3400 and NZ3400*, were used for the immobilized and planktonic adaptation, respectively. Occurrence of identical mutants recovered from immobilized and planktonic adaptation with the SNP in LP_RS14990 seems therefore unlikely to originate from a stock subculture, as confirmed by Sanger sequencing (data not shown). Furthermore, the mutation was not detected by Pyrosequencing of NZ3400* before inoculation (data not shown), minimizing a role for subcultures in our experiments. Moreover, the increased fitness of mutants shows the beneficial effect of detected mutations. This strongly supports that observed mutations are caused by adaptive evolution and demonstrates the suitability of the PolyFermS model to select relevant mutations related to the human gut microbiota. The exact nature of the evolutionary pressure in our system is not known and remains to be elucidated. It is likely the result of a combination of factors such as the competition for nutrients, metabolic cross-feeding, physiochemical factors, or presence of antimicrobial metabolites.

In conclusion, we demonstrated successful application of the continuous PolyFermS gut fermentation model to provide a long-term stable gut microbiota to generate adapted mutants to this environment. Immobilization of strains does not only allow adaptive evolution of non-colonizers but also creates a culture consisting of sessile and planktonic cells mimicking the human gastrointestinal tract. The conditions of the model can be easily changed to other needs and selective pressures, including the source of microbiota and abiotic conditions. This novel technology enables identification of genes involved in gut microbiota colonization, persistence, and metabolism. The PolyFermS system could further be designed and applied for different fermentations, to trace and identify evolutionary and ecological processes between an exogenous single strain and a complex ecosystem.

## Acknowledgment

We thank Alfonso Die for assistance during HPLC measurements, Dr. Florentin Constancias for his support converting 16S sequencing raw data into amplicon sequence variants, and Julia Iselin for experimental assistance. Pyrosequencing was set-up in collaboration with the Genetic Diversity Centre (GDC), ETH Zurich. The study was funded by the ETH Zurich research grant program (ETH-30 15-2).

## Caption Supplemental Figures

**Fig. S1:** Comparison of metabolite concentrations in EMS medium and the reactor effluent. Barplot representing metabolite concentrations (mM) of the EMS medium consisting of reactor effluent, MacFarlane medium and glucose and the reactor effluent without *L. plantarum* supplementation. Samples were analyzed in biological triplicates.

**Fig S2:** Concentrations (mM) of main SCFAs of PolyFermS effluent samples inoculated with immobilized and planktonic *L. plantarum*: A) IR1 inoculated with donor 1 immobilized fecal microbiota and immobilized *L. plantarum*; B) IR2 inoculated with donor 2 immobilized fecal microbiota and connected treatment reactors (TRs) C) TR1 (period 1), D) TR1 (period 2)and E) TR2 spiked with planktonic *L. plantarum*.

**Fig. S3:** Microbial composition in relative abundance obtained by 16S rRNA amplicon sequencing. Composition at phylum (A,C) and family (B,D) level of *in vitro* proximal colon microbiota of A,B) IR1 inoculated with donor 1 immobilized fecal microbiota and immobilized *L. plantarum* and of C,D) IR2 inoculated with donor 2 immobilized fecal microbiota and connected treatment reactors (TRs) TR1 (period 1), TR1 (period 2) and TR2 spiked with planktonic *L. plantarum*. Values at family level of < 1% are summarized in the “Others” group. X-axis labeling indicates the day during fermentation relative to the start of the IR.

**Fig. S4:** Principal component analysis (PCA) of growth pattern of recovered *L. plantarum* strains. *L. plantarum* strains were recovered from the immobilized and planktonic adaptation cycle and grown in MRS and MRS supplemented with acetate (50 mM), propionate (30 mM) and butyrate (30 mM). Reference indicates the reference strain NZ3400*. Plot is based on the R function fviz_pca_ind.

**Fig. S5:** Growth of *L. plantarum* in MRS during four weeks daily serial cultivation. *L. plantarum* was serially cultivated in triplicates in MRS medium for four weeks. Final OD was measured in the first overnight culture (t0) and after one, two, three and four weeks (t1-t4). ΔOD600 values were calculated by subtracting values obtained after 16 h of growth at 37°C by values obtained at the time point of inoculation. Values represent mean ± standard deviation of triplicates. The reference strain (RT) and selected recovered *L. plantarum* strains from immobilized and planktonic adaptation are depicted on the x-axis.

**Fig. S6:** Pyrogram obtained by Pyrosequencing of the C837A SNP in LP_RS15205. A) Pyrogram obtained from donor 2 gut microbiota prior to *L. plantarum* supplementation. B) Pyrogram obtained from donor 3 gut microbiota supplemented with *L. plantarum* PA2_06 and NZ3400* in a ratio 1:1. X-axis depicts the added nucleotide over time and y-axis represents the light signal induced by incorporated nucleotides. Allele frequency was measured for G and T since Pyrosequencing was done on the complementary DNA strand.

## Caption Supplemental Tables

**Table S1:** Strains and plasmids used in this study.

**Table S2:** Primers used in this study.

**Table S3:** SNP stability in *L. plantarum* IA10 and PA2_06 during 12 days of continuous cultivation in MRS medium.

## Supplemental Material and Methods

Supplementary Data S1

