## Supplementary Material for "*In vitro* gut modeling as a tool for adaptive evolutionary engineering of *Lactiplantibacillus plantarum*"

Supplementary Data 1 – Methods selection

### **MacFarlane medium composition and supplemented vitamins**

| **Component** | **g/l** | **Vitamin** | **µg/l** |
| --- | --- | --- | --- |
| Pectin (citrus)  Yeast extract  Xylan (oat spelts)  Arabinogalactan (larch wood)  Guar gum  Inulin (Raftiline® HP)  Soluble starch (potato)  Mucin  Casein acid hydrolysate  Peptone water  Bacto^TM^ Tryptone  L-Cysteine HCl  Bile Salts  KH_2_PO_4_  NaHCO_3_  NaCl  KCl  MgSO_4_ anh. (M: 120.37)  CaCl_2_*2H_2_O (M: 147.02)  MnCL_2_*4H_2_O (M: 197.91)  FeSO_4_*7H_2_O (M: 278.02)  Hemin solution  Tween 80  Vitamin solution | 2  4.5  2  2  1  1  5  4  3  5  5  0.8  0.4  0.5  1.5  4.5  4.5  0.61  0.1  0.2  0.005  1 ml  1 ml  1 ml | Pyridoxine-HCl (Vit. B_6_)  4-Aminobenzoic acid (PABA)  Nicotinic acid (Vit. B_3_)  Biotin (Vit. H), stock: 4mg/ml  Folic acid (Vit. B_9_), stock: 4mg/ml  Cyanocobalamine (Vit. B_12_)  Thiamine (Vit. B_1_HCl)  Riboflavine (Vit. B_2_)  Phylloquinone (Vit. K_1_), stock: 0.15 mg/ml  Menadione (Vit. K_3_), stock: 2 mg/ml  Pantothenate (Vit. B_5_) | 100  50  50  20  20  5  50  50  0.075  10  100 |
| Adjusted pH to 5.8  Filter-sterilized vitamin solution added after autoclaving | | | |

### **Microbial profiling by 16S rRNA gene amplicon sequencing**

Removal of Illumina adaptors and gene-specific primers was done using Cutadapt (1). Sequences were processed using the DADA2 pipeline (2) which allows inference of exact amplicon sequence variants (ASVs). Forward and reverse reads were truncated after 170 and 160 nucleotides, respectively. After truncation, reads with expected error rates higher than three and four for forward and reverse reads, respectively, were removed. After filtering, error rate learning, ASV inference, and denoising, reads were merged with a minimum overlap of 40 bp. Chimeric sequences were identified and removed. Taxonomy was assigned using the DADA2 formatted SILVA database (v.132) (3).

### **Competition experiments in the human gut microbiota**

Primers (Supplementary Table 2) selectively targeting *L. plantarum* in the gut microbiota up- and downstream of investigated SNPs and sequencing primers were designed as described previously (4) using PyroMark^TM^ID 1.0 software (Biotage AB and Biosystems, Uppsala, Sweden). To determine the background of the method, DNA isolated from gut microbiota before *L. plantarum* addition was screened. If an allele was absent in the gut microbiota, whole microbiota DNA was used. In case of background, exogenously added *L. plantarum* was enriched by adding 100 μl reactor effluent to 10 ml MRS + CM and grown at 37°C overnight. Cells were harvested and DNA was isolated using the FastDNA® SPIN Kit for Soil.

Biotin amplification by PCR was performed using the 2X PCR Master Mix (Life Technologies Europe BV, Zug, Switzerland) and 40 ng DNA in a T3000 Thermocycler (Biometra, Göttingen, Germany) as described previously (4). Immobilization on streptavidin-coated beads (GE Healthcare Bio-Sciences AB, Uppsala, Sweden), hybridization of the sequencing primer to the DNA and product washing was done as described previously using a PyroMark^TM^ Vacuum Prep Worktable (Biotage, Uppsala, Sweden) (5). Pyrosequencing was performed at the Genetic Diversity Centre (Zürich, Switzerland) using a PyroMark Q96 ID (Biotage, Uppsala, Sweden) system.

### **Plasmid construction and gene replacement of *L. plantarum* NZ3400* LP_RS14990 gene**

The knockout plasmid was generated *in silico* using a cassette encoding an rRNA adenine N-6-methyltransferase erythromycin resistance gene (*ery*) (6) flanked by the 1050 bp up- and downstream sequence of the LP_RS14990 gene with 50 bp overlap with the LP_RS14990. This sequence was synthesized at Biocat and copied into the pUC18 vector (Biocat, Heidelberg, Germany), resulting in the LP_RS14990 gene replacement vector pUC18_lp_lamC. The vector was transferred into calcium competent *E. coli* MC1000 (7). *E. coli* MC1000 was grown overnight in Brain Heart Infusion Broth (BHI, Labolife Sàrl, Pully, Switzerland) at 37°C with agitation. The vector was then isolated from full-grown cultures using the PureLink® Quick Plasmid Miniprep Kit (Invitrogen, Thermo Fisher Scientific Inc., Waltham, USA). The purified plasmid was transferred to *L. plantarum* NZ3400* by electroporation using an Eppendorf Eporator® (Eppendorf, Hamburg, Germany) as described previously (8). Plasmid integration was detected by plating on MRS agar + 50 µg/ml erythromycin. Double cross-over strains in which LP_RS14990 was replaced by *ery,* were identified by PCR using specific primers (see Table S2 in the supplemental material) targeting the LP_RS14990 gene (lamC-5’, lamC-3’) and primers binding outside the flanking region cloned in pUC18_lp_lamC vector (lamC-D-5’, lamC-U-3’) and the erythromycin gene (ery-5’, ery-3’).

Supplementary Table S1: Strains and plasmids used in this study.

|  | **Relevant features** | **Source of reference** |
| --- | --- | --- |
| ***L. plantarum*** |  |  |
| NZ3400 | Reference strain, WCFS1 derivative containing a lox66-P32-cat-lox71 insertion in the neutral H-locus | Remus *et al.*, 2012 |
| IA_10.2 | Derivative strain of IA10 with C837A in LP_RS15205 and G984A in LP_RS10985, day 23, Cm^R^ | This study |
| IA_10.3 | Derivative strain of IA10 with C569A in LP_RS14255, day 23, Cm^R^ | This study |
| PA1.1_06 | Strain from biofilm of planktonic adaptation, TR1 (period1), day 44, Cm^R^ | This study |
| PA1.1_07 | Strain from biofilm of planktonic adaptation, TR1 (period1), day 44, Cm^R^ | This study |
| PA1.2_01 | Strain with C979T in LP_RS14990 and G382A in LP_RS01530 from biofilm of planktonic adaptation, TR1 (period2), day 23, Cm^R^ | This study |
| PA1.2_02 | Strain with C979T in LP_RS14990*,* G382A in LP_RS01530 and intergenic SNP LP_RS05100 < LP_RS05095 from biofilm of  planktonic adaptation, TR1 (period2), day 23, Cm^R^ | This study |
| PA1.2_03 | Strain with C979T in LP_RS14990*,* G382A in LP_RS01530 and C2340T in LP_RS04385 from biofilm of planktonic adaptation,  TR1 (period2), day 23, Cm^R^ | This study |
| PA2_03 | Strain with C749T in LP_RS15260 from biofilm of planktonic adaptation, TR2, day 72, Cm^R^ | This study |
| PA2_07 | Strain with A767G in LP_RS05980 from biofilm of planktonic adaptation, TR2, day 72, Cm^R^ | This study |
| PA2_08 | Strain with C995T in LP_RS01370 and G881A in LP_RS07205 from biofilm of planktonic adaptation, TR2, day 72, Cm^R^ | This study |
| PA2_13 | Strain from biofilm of planktonic adaptation, TR2, day 72, Cm^R^ | This study |
| PA2_14 | Strain from biofilm of planktonic adaptation, TR2, day 72, Cm^R^ | This study |
| PA2_15 | Strain from biofilm of planktonic adaptation, TR2, day 72, Cm^R^ | This study |
| IA_10.1 | Derivative strain of IA10 with C569A in LP_RS14255, day 23, Cm^R^ | This study |
| IA_10.4 | Derivative strain of IA10 with C569A in LP_RS14255, day 23, Cm^R^ | This study |
| IA_10.5 | Derivative strain of IA10 with C569A in LP_RS14255, day 23, Cm^R^ | This study |
| PA1.1_01 | Strain with G173A in LP_RS08140 from planktonic adaptation, TR1 (period1), day 44, Cm^R^ | This study |
| PA1.1_02 | Strain with G645T in LP_RS06730 and G1192A in LP_RS02260 from planktonic adaptation, TR1 (period1), day 44, Cm^R^ | This study |
| PA1.1_03 | Strain with A487C in LP_RS12455 and C328T in LP_RS13325 from planktonic adaptation, TR1 (period1), day 44, Cm^R^ | This study |
| PA1.1_04 | Strain from planktonic adaptation, TR1 (period1), day 41, Cm^R^ | This study |
| PA1.1_05 | Strain from planktonic adaptation, TR1 (period1), day 41, Cm^R^ | This study |
| PA2_01 | Strain with intergenic SNP LP_RS05100 < LP_RS05095 and C749T in LP_RS15260 from planktonic adaptation, TR2, day 64, Cm^R^ | This study |
| PA2_02 | Strain with C749T in LP_RS15260 from planktonic adaptation, TR2, day 72, Cm^R^ | This study |
| PA2_04 | Strain with C837A in LP_RS15205 from planktonic adaptation, TR2, day 72, Cm^R^ | This study |
| PA2_05 | Strain with C837A in LP_RS15205 from planktonic adaptation, TR2, day 72, Cm^R^ | This study |
| PA2_06 | Strain with C837A in LP_RS15205 from planktonic adaptation, TR2, day 72, Cm^R^ | This study |
| PA2_09 | Strain from planktonic adaptation, TR2, day 64, Cm^R^ | This study |
| PA2_10 | Strain from planktonic adaptation, TR2, day 72, Cm^R^ | This study |
| PA2_11 | Strain from planktonic adaptation, TR2, day 72, Cm^R^ | This study |
| PA2_12 | Strain from planktonic adaptation, TR2, day 72, Cm^R^ | This study |

Supplementary Table S1, continued

|  | **Relevant features** | **Source of reference** |
| --- | --- | --- |
| ***L. plantarum*** |  |  |
| IA01 | Strain with C979T in LP_RS14990 from immobilized adaptation, day 53, Cm^R^ | This study |
| IA02 | Strain with C979T in LP_RS14990 from immobilized adaptation, day 53, Cm^R^ | This study |
| IA03 | Strain with C979T in LP_RS14990 and G39T in LP_RS13860 from immobilized adaptation, day 53, Cm^R^ | This study |
| IA04 | Strain with C569A in LP_RS14255 from immobilized adaptation, day 53, Cm^R^ | This study |
| IA05 | Strain with C569A in LP_RS14255 from immobilized adaptation, day 53, Cm^R^ | This study |
| IA06 | Strain with C569A in LP_RS14255 from immobilized adaptation, day 53, Cm^R^ | This study |
| IA07 | Strain with C569A inLP_RS14255 from immobilized adaptation, day 53, Cm^R^ | This study |
| IA08 | Strain with C569A in LP_RS14255 from immobilized adaptation, day 53, Cm^R^ | This study |
| IA09 | Strain with C569A in LP_RS14255 from immobilized adaptation, day 53, Cm^R^ | This study |
| IA10 | Strain with C569A in LP_RS14255 from immobilized adaptation, day 53, Cm^R^ | This study |
| IA11 | Strain with intergenic SNP LP_RS00275 < LP_RS00270 from immobilized adaptation, day 53, CmR | This study |
| IA12 | Strain from immobilized adaptation, day 53, Cm^R^ | This study |
| Strain1-7 | Strains from planktonic adaptation, used to assess phenotypic stability |  |
| ***E. coli*** |  |  |
| MC1000 |  | Casadaban *et al*., 1980 |
| ***Plasmid*** |  |  |
| pUC18_lp_  lamC | Em^R^, gene replacement vector | This study |

Cm^R^: chloramphenicol-resistance, Em^R^: erythromycin-resistance

Supplementary Table S2: Primers used in this study.

| **Name** | **Sequence (5'-3')** | **Description** |
| --- | --- | --- |
| lamC-5' | TGAGATACTCGCTTCACCGTC | Targeting LP_RS14990 |
| lamC-3' | ATGAGTTGCGACGTTTCAGAC | Targeting LP_RS14990 |
| lamC-D-5' | CGTCTACACCATGTTCGGTTA | Targeting flanking region of LP_RS14990 |
| lamC-U-3' | GAAGGCAATTCGGCGTAAC | Targeting flanking region of LP_RS14990 |
| ery-5' | GAATCGAGACTTGAGTGTG | Targeting *ery* gene |
| ery-3' | CACACTCAAGTCTCGATTC | Targeting *ery* gene |
| lamC-Pyro-D-5' | GGGTGGTAACTGTTCAAAAACATT | Pyrosequencing, downstream of the SNP C979T in LP_RS14990 |
| lamC-Pyro-U-3' | AGCTATGCACAATCGCAAGAGA | Pyrosequencing, upstream of the SNP C979T in LP_RS14990 |
| lamC-seq | TGTCCTACCGATGCTT | Pyrosequencing, sequencing primer LP_RS14990 |
| 3630-Pyro-D-5' | TGGGATTGTTCAAAATCAACTG | Pyrosequencing, downstream of the SNP C979T in LP_RS15205 |
| 3630-Pyro-U-3' | AGGTCAAGATTGCCACGTTAA | Pyrosequencing, upstream of the SNP C979T in LP_RS15205 |
| 3630-seq | CCGATGAAGCGAACT | Pyrosequencing, sequencing primer LP_RS15205 |

Supplementary Table S3: SNP stability in *L. plantarum* IA10 and PA2_06 during 12 days of continuous cultivation in MRS medium.

|  | Allele frequency (%) at day 0 | | | Allele frequency (%) at day 12 | | |
| --- | --- | --- | --- | --- | --- | --- |
|  | Replicate 1 | Replicate 2 | Replicate 3 | Replicate 1 | Replicate 2 | Replicate 3 |
| LP_RS14990gene ^a)^ |  |  |  |  |  |  |
| C | 0 | 0 | 0 | 0 | 0 | 0 |
| T | 100 | 100 | 100 | 100 | 100 | 100 |
| LP_RS15205 gene^b)^ |  |  |  |  |  |  |
| G | 0 | 0 | 0 | 3.6 | 3.3 | 3.5 |
| T | 100 | 100 | 100 | 96.4 | 96.7 | 96.5 |

a) Allele frequency of the LP_RS14990 mutation C979T in *L. plantarum* IA01 culture.

b) Allele frequency of the LP_RS15205 mutation C837A in *L. plantarum* PA2_06 culture. Allele frequency was measured for G and T since Pyrosequencing was done on the complementary DNA strand.

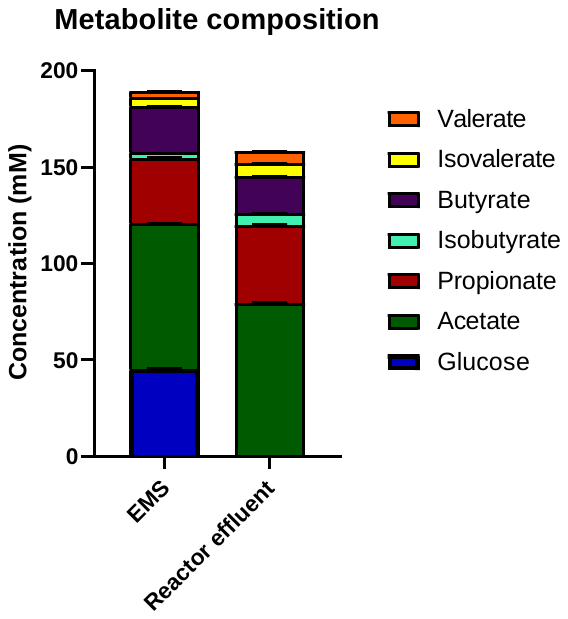

**Supplementary Figure S1: Comparison of metabolite concentrations in EMS medium and the reactor effluent.** Barplot representing metabolite concentrations (mM) of the EMS medium consisting of reactor effluent, MacFarlane medium and glucose and the reactor effluent without *L. plantarum* supplementation. Samples were analyzed in biological triplicates.

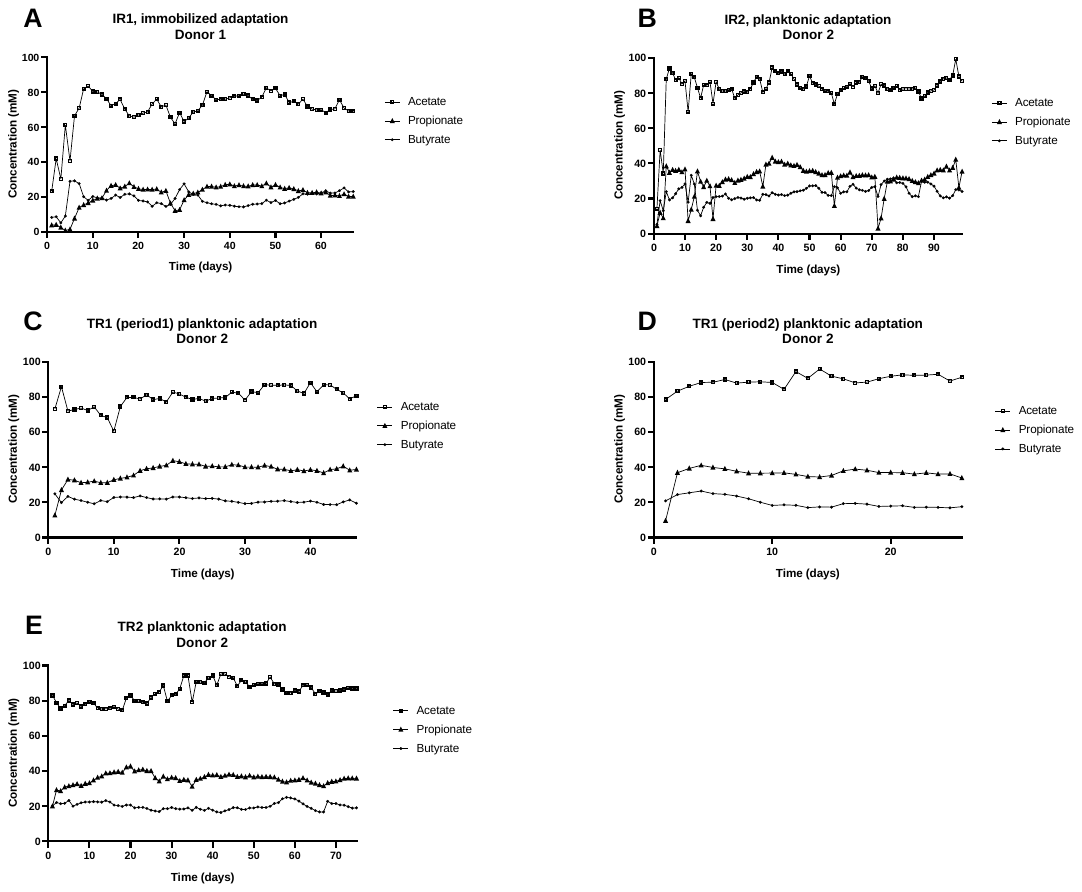

**Supplementary Figure S2:** Concentrations (mM) of main SCFAs of PolyFermS effluent samples inoculated with immobilized and planktonic *L. plantarum*: A) IR1 inoculated with donor 1 immobilized fecal microbiota and immobilized *L. plantarum*; B) IR2 inoculated with donor 2 immobilized fecal microbiota and connected treatment reactors (TRs) C) TR1 (period 1), D) TR1 (period 2)and E) TR2 spiked with planktonic *L. plantarum*.

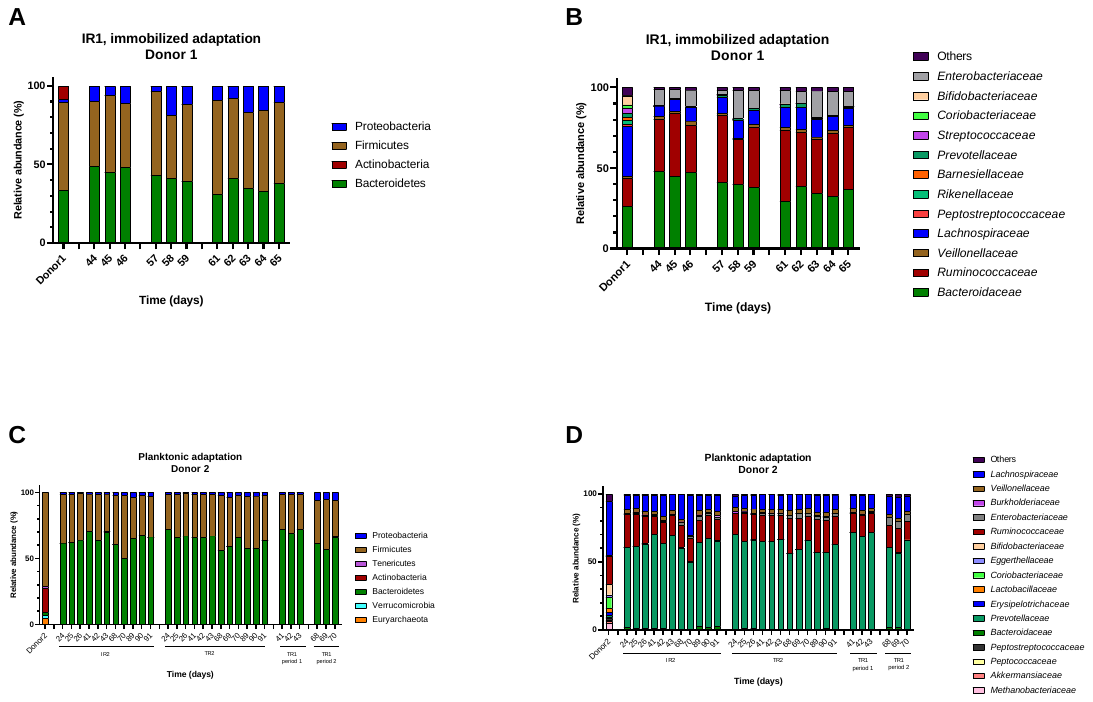

**Supplementary Figure S3:** Microbial composition in relative abundance obtained by 16S rRNA amplicon sequencing. Composition at phylum (A,C) and family (B,D) level of *in vitro* proximal colon microbiota of A,B) IR1 inoculated with donor 1 immobilized fecal microbiota and immobilized *L. plantarum* and of C,D) IR2 inoculated with donor 2 immobilized fecal microbiota and connected treatment reactors (TRs) TR1 (period 1), TR1 (period 2) and TR2 spiked with planktonic *L. plantarum*. Values at family level of < 1% are summarized in the “Others” group. X-axis labeling indicates the day during fermentation relative to the start of the IR.

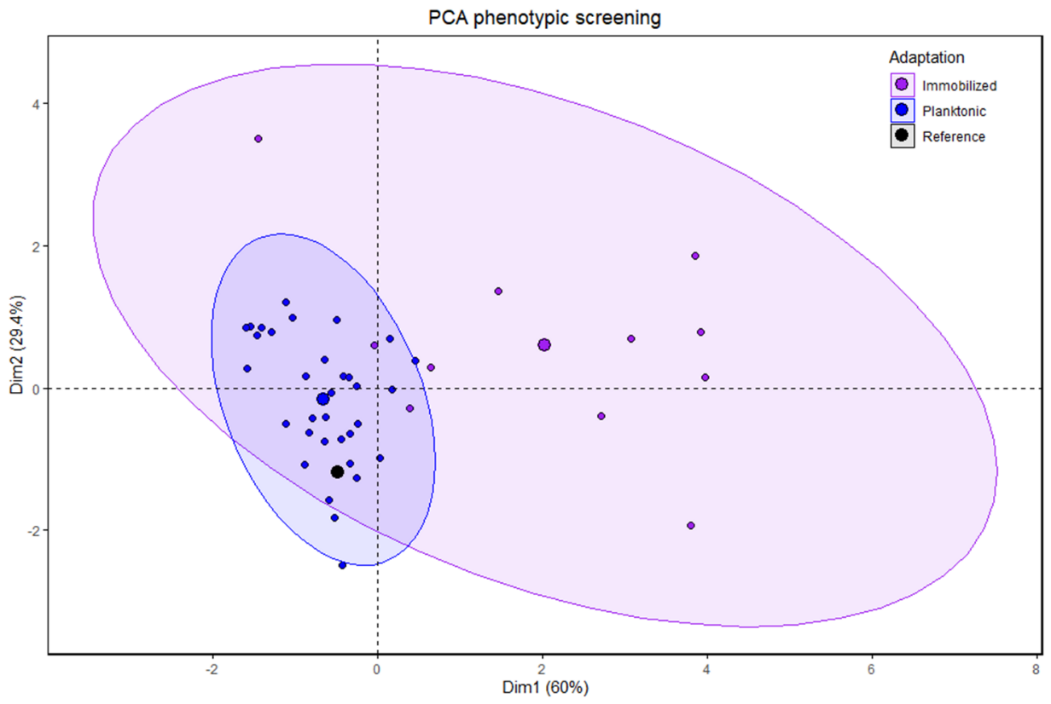

**Supplementary Figure S4: Principal component analysis (PCA) of growth pattern of recovered *L. plantarum* strains.** *L. plantarum* strains were recovered from the immobilized and planktonic adaptation cycle and grown in MRS and MRS supplemented with acetate (50 mM), propionate (30 mM) and butyrate (30 mM). Reference indicates the reference strain NZ3400*. Plot is based on the R function fviz_pca_ind.

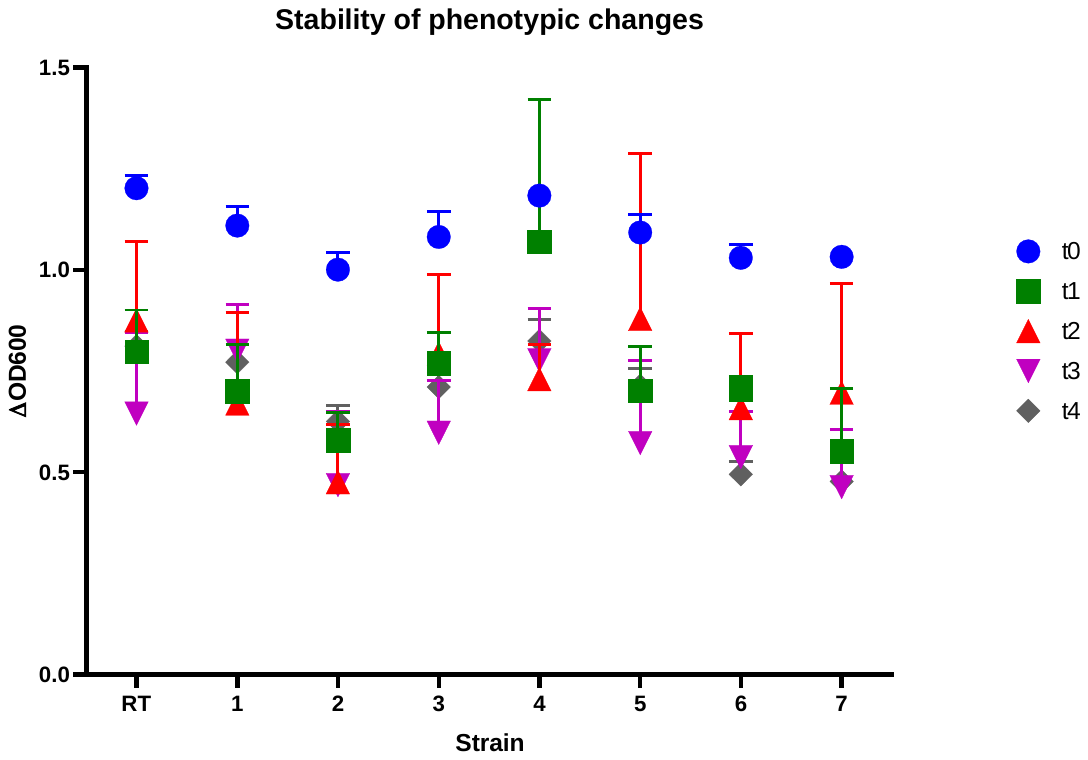

**Supplementary Figure S5: Growth of *L. plantarum* in MRS during four weeks daily serial cultivation.** *L. plantarum* was serially cultivated in triplicates in MRS medium for four weeks. Final OD was measured in the first overnight culture (t0) and after one, two, three and four weeks (t1-t4). ΔOD600 values were calculated by subtracting values obtained after 16 h of growth at 37°C by values obtained at the time point of inoculation. Values represent mean±standard deviation of triplicates. The reference strain (RT) and selected recovered *L. plantarum* strains from immobilized and planktonic adaptation are depicted on the x-axis.

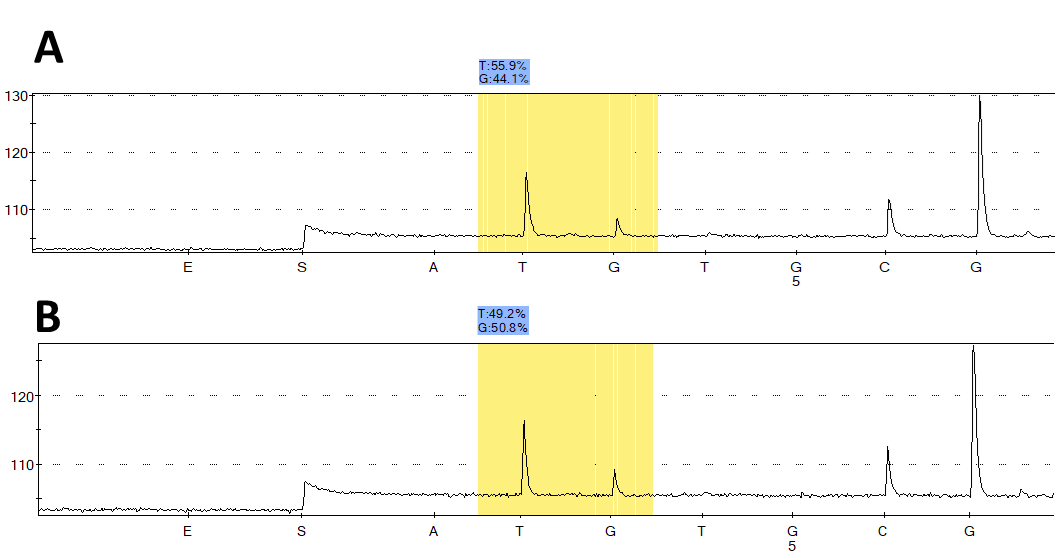

**Supplementary Figure S6: Pyrogram obtained by Pyrosequencing of the C837A SNP in LP_RS15205.** A) Pyrogram obtained from donor 2 gut microbiota prior to *L. plantarum* supplementation. B) Pyrogram obtained from donor 3 gut microbiota supplemented with *L. plantarum* PA2_06 and NZ3400* in a ratio 1:1. X-axis depicts the added nucleotide over time and y-axis represents the light signal induced by incorporated nucleotides. Allele frequency was measured for G and T since Pyrosequencing was done on the complementary DNA strand.
